# Microbial Source Tracking in the Love Creek Watershed, Delaware (USA)

**DOI:** 10.1101/2020.01.13.900647

**Authors:** Christopher R. Main, Robin Tyler, Karen Lopez, Serigo Heurta

## Abstract

Fecal contamination of waterways in Delaware pose an ongoing problem for environmental and public health. For monitoring efforts, *Enterococcus* has been widely adopted by the state to indicate the presence of fecal matter from warm-blooded animals and to establish Primary and Secondary Contact Recreation criteria. In this study, we examined sites within the Love Creek watershed, a tributary of the Rehoboth bay, using next-generation sequencing and SourceTracker to determine sources of potential fecal contamination and compared to bacterial communities to chemical and nutrient concentrations. Microbial community from fecal samples of 10 different types of animals and 1 human sample were used to generate a fecal library for community-based microbial source tracking. Orthophosphate and total dissolved solids were among the major factors associated with community composition. SourceTracker analysis of the monthly samples from the Love Creek watershed indicated the majority of the microbial community were attributed to “unknown” sources, i.e. wildlife. Those that attribute to known sources were primarily domestic animals, i.e. cat and dog. These results suggest that at the state level these methods are capable of giving the start for source tracking as a means to understanding bacterial contamination.

## Introduction

The Department of Natural Resources and Environmental Control (DNREC) Environmental Laboratory Section (ELS) has been monitoring the waters of Delaware for several decades as a requirement of the Clean Water Act (CWA) (USEPA 1987 - Sections 106, 303, 304 and 305). Section 303(d) identifies “impaired” waters as those that do not meet the Water Quality Standards laid out in Section 304(a). It includes processes for determining the degree of pollution reduction from human related sources likely to result in attainment of the Total Maximum Daily Load (TMDL) and recommending and informing land and water management practices necessary to achieve the targeted pollution reduction. Routine long-term monitoring at established stations tracks water quality status and documents how State waters are responding to environmental stewardship efforts.

One measurable type of potential human-related water pollution is fecal bacteria. These bacteria may derive from humans, various domesticated animals, i.e. dogs, cats, cows, etc. or from wild animals. Bacteria within the genus *Enterococcus* has been widely adopted to indicate the presence of fecal matter from warm-blooded animals in 305(b) monitoring efforts. Delaware has risk-based numeric criteria for Primary Contact Recreation (PCR) and Secondary Contact Recreation (SCR) in freshwater and saltwater “determined by the Department (DNREC) to be of non-wildlife origin based on best scientific judgment using available information” (DNREC, 2014a). Important among the limitations of the *Enterococcus* test is that it does not differentiate as to what type of animal the detected bacteria are from. It could be any mix of warm-blooded animal types that might be in the watershed – i.e. mammals and birds, domestic and wildlife.

While these *Enterococcus* criteria are generally recognized to be protective of human health and the continuation of routine monitoring of waters is necessary to track status and trends, this test does not help in locating sources of bacterial contamination, which may be abatable. A supplemental, complimentary addition to the existing monitoring format is needed to facilitate where to apply pollution control practices and maximize the frugal utilization of the increasingly scarce resources available to bring about the environmental improvements intended under the CWA, such that waters meet standards criteria and attain designated uses (DNREC, 2014a), for example, PCR, SCR, fish, aquatic life and wildlife and harvestable shellfish waters.

Delaware’s inland coastal bays (DIB) consist of three interconnected water bodies, Rehoboth, Indian River and Little Assawoman bays that drain approximately 300 square miles of mixed land use. Eutrophication of the DIB has increased over the last several decades with inputs from agricultural and urban sources (Price, 1998; Sallade and Sims, 1997). The Love Creek watershed (Fig. 1) is a tributary of the Rehoboth bay and part of the National Estuary Program for over 20 years. The watershed has undergone extensive human development of various types in its’ tidal and non-tidal segments with the inevitability of substantially more, making for an ideal study site. Its’ environmental condition, aquatic and terrestrial, could benefit substantially from improved precision in identifying human-related pollution sources. Additionally, considerable periodic seabird activity in the tidal segment and relatively wide forested stream corridors in the non-tidal segment are evidence of a robust complement of indigenous wildlife, e.g. deer, raccoons.

**Figure 1.**
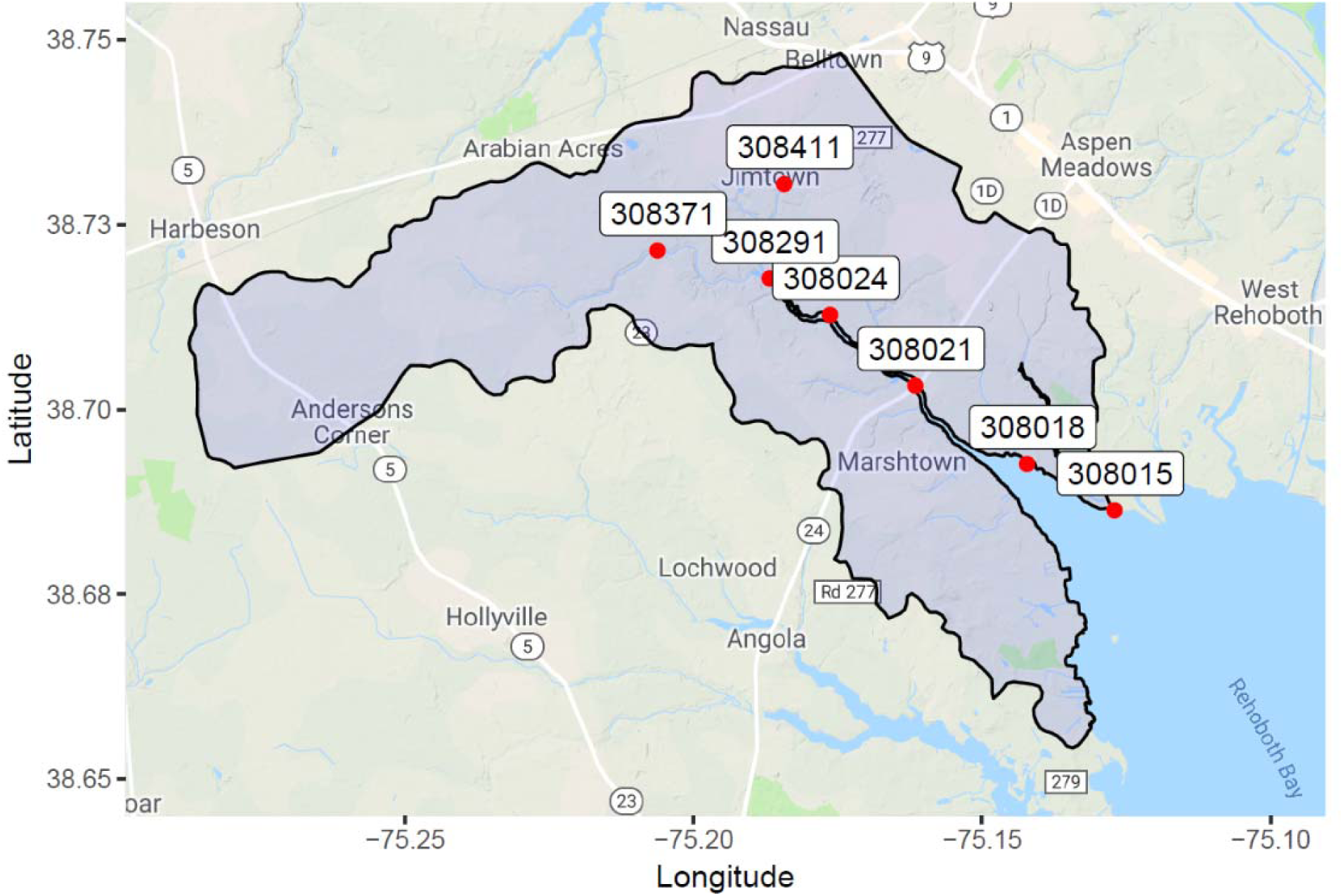
Sampling sites in the Love Creek Watershed. See Table 1 for more information.

Microbial source tracking (MST) methods use specific bacterial profiles associated with hosts, e.g. human, farm animal, bird, to determine sources of fecal contamination in the environment. The most common MST method uses quantitative real-time polymerase chain reaction (qPCR) and primers that target the 16S rRNA genes of host associated bacteria (Brown et al., 2017; Harwood et al., 2014), however, qPCR may suffer from specificity and sensitivity issues (Brown et al., 2019; Green et al., 2014; Stewart et al., 2013). In the last decade, advancements in high-throughput DNA sequencing has led to large-scale microbial community studies. Community based MST analysis is generated from unique microbial community profiles of environmental and fecal sources leveraged from next generation sequencing (NGS) techniques (Unno et al., 2018). Two methods may be used to define the microbial community: 1) assignment of operational taxonomic units (OTUs) via a clustering algorithm, usually to a 97% similarity between sequences, or 2) amplicon sequence variant (ASV) which can resolve difference in gene regions to a single nucleotide (Callahan et al., 2017).

**Table 1:**
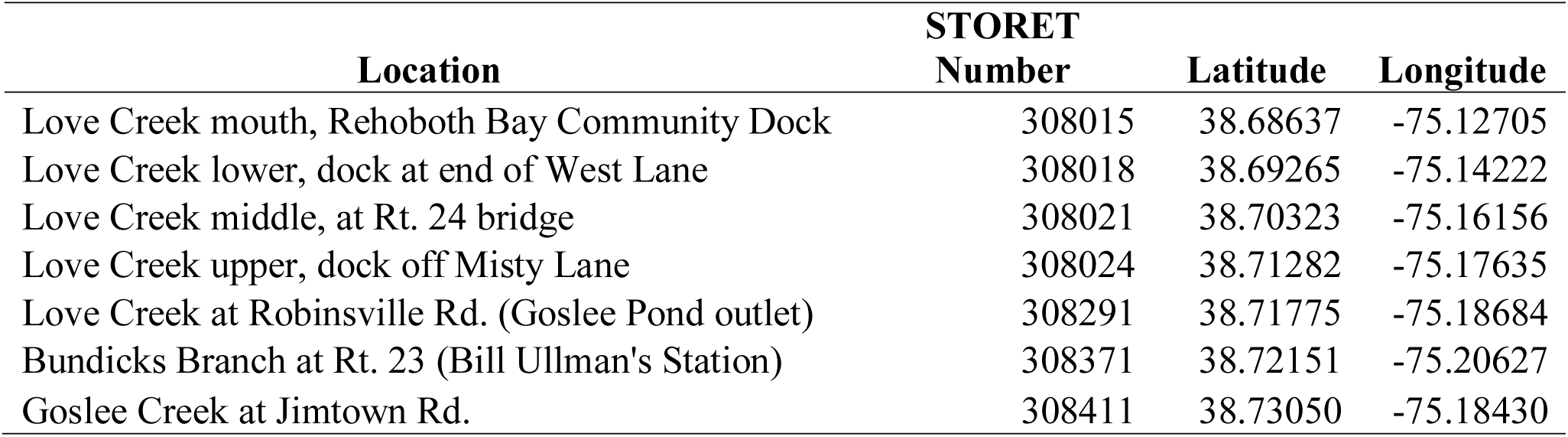
Location Descriptions, STORET Numbers and Latitude and Longitude of Sampling Sites.

| **Location** | **STORET**<br>**Number** | **Latitude** | **Longitude** |
| --- | --- | --- | --- |
| Love Creek mouth, Rehoboth Bay Community Dock | 308015 | 38.68637 | -75.12705 |
| Love Creek lower, dock at end of West Lane | 308018 | 38.69265 | -75.14222 |
| Love Creek middle, at Rt. 24 bridge | 308021 | 38.70323 | -75.16156 |
| Love Creek upper, dock off Misty Lane | 308024 | 38.71282 | -75.17635 |
| Love Creek at Robinsville Rd. (Goslee Pond outlet) | 308291 | 38.71775 | -75.18684 |
| Bundicks Branch at Rt. 23 (Bill Ullman's Station) | 308371 | 38.72151 | -75.20627 |
| Goslee Creek at Jimtown Rd. | 308411 | 38.73050 | -75.18430 |

SourceTracker is a Bayesian classification program, which provides an estimated percentage of the sequenced microbial community from an environmental samples can be attributed to a specific fecal source (Brown et al., 2019; Knights et al., 2011). SourceTracker uses a Gibbs sampling algorithm to examine the likely prior distribution of OTUs/ASVs within user-defined sources. This distribution within the sources can be used to determine the affiliation within sample (referred to as sinks) communities and the contribution of each source is then determined based on this distribution. The SourceTracker algorithm has been shown to be more accurate than random forest analysis, or naive Bayesian classification (Knights et al., 2011). SourceTracker has been used to determine contamination sources in the Russian River (Dubinsky et al., 2016), ATM keypads (Bik et al., 2016), recreational beaches in Australia (Ahmed et al., 2015; Brown et al., 2017; Henry et al., 2016; McCarthy et al., 2017), lakes in St. Paul, Minnesota (Brown et al., 2019), and the upper Mississippi River (Staley et al., 2015, 2014).

Our goals were to use next generation sequencing to create community based microbial profiles of fecal sources and monthly samples of sites within the Love Creek watershed. Using SourceTracker we examined the bacterial inputs from various fecal sources into the monthly samples. In addition, we examined the impact of a significant rain event, within 24 hours, at the tidally driven sites. The results of this pilot study are broadly relevant to assessing fecal pollution within the waterways of Delaware.

## Material and Methods

### Study Site and Sample Collection

The Love Creek watershed is drains approximately 24 square miles of land into the Rehoboth Bay (Fig. 1) and is tidally driven to the dam at Goslee Pond (STORET: 308291; Homsey et al., 2015). In the last two decades urban development has increased approximately 80% with a loss of forested uplands and agriculture (Homsey et al., 2015). Water samples were collected monthly from March to October 2017 from seven sites in the Love Creek watershed (Table 1), with triplicate samples collected at the Rt. 24 Marina (STORET: 308021). Sites were considered marine waters when salinity was greater than 5 (DNREC, 2014a, pg. 3). To examine the influence of rainfall on microbial transport, rainfall was monitored using the Delaware Environmental Observing System at the Millsboro Long Neck station (DLNK; http://deos.udel.edu/odd-divas/station_daily.php?network=DEOS&station=DLNK). Samples at the tidally driven sites (STORET: 308015, 308018, 308021, and 308024) were collected 24 hours after a rainfall event of approximately 62 mm of rain in 2 hours. Temperature, salinity, dissolved oxygen (mg l^−1^), pH, and specific conductivity were measured using a YSI 650 (YSI Inc., Yellow Springs, OH). Dissolved nutrients (NO_3_ plus NO_2_ [NO_X_], OP, organic carbon, and dissolved solids), and total nutrients (chlorophyll *a*, nitrogen, phosphorus, organic carbon, turbidity and suspended solids) concentrations were determined by APHA Standard Methods (American Public Health Association, 2012). *Enterococcus* concentrations were determined for each sample using Enterolert© (IDEXX, Westbrook, ME).

### Digital Elevation Analysis

High-resolution 2014 LiDAR data was used to identify potential portions of the landscape that were high potential for producing fecal contamination. The 1-m LiDAR-based digital elevation model (DEM) was smoothed using focal statistics with a 7 by 7 filter to remove microtopography. ArcGIS (ESRI, 2017) and TauDEM (Tarboton, 1997) software were used to analyze the DEM, create a hydro-enforced DEM, identify topographically convergent areas, flow pathways, and hydrologic connectivity. This topographic connectivity model was used in conjunction with information about the locations of residential septic systems (DNREC, 2014b), soil hydrologic groups (NRCS, 2016), and hydrogeology (Andres, 2004; Martin and Andres, 2008), to identify hydrologically sensitive areas (Agnew et al., 2006; Walter et al., 2000), areas where bacteria or nutrient loading may have a more direct contribution to downstream contamination.

### Sample Filtering and Processing

Water samples were filtered within 6 hours of sampling under gentle vacuum (∼380 mm Hg) on 0.45-µm polycarbonate filters (Millipore Isopore, Billerica, MA). Filters were immediately placed into CTAB buffer (100 mM Tris-HCl (pH 8), 1.4 M NaCl, 2% (wt/vol) cetyltrimethylammonium bromide (CTAB), 0.4% (vol/vol) 2-mercaptoethanol, 1% (wt/vol) polyvinylpyrrolidone, and 20 mM EDTA; Dempster et al., 1999) and stored at −80°C until extraction. Fecal samples were collected in July 2016 during the Delaware State Fair. Bovine, chicken, goat, horse, pig, sheep, domesticated duck and goose samples were collected with as much metadata information as possible, i.e. location, feed, sex, etc. by Christopher Main and Karen Lopez. Cat, dog and human samples were collected by or from Christopher Main and all samples were stored at −80°C until extraction.

Before extraction, filters were heated at 65°C for 10 minutes. Following incubation, 700 µl of a 24:1 isoamyl:chloroform solution was added to each sample and briefly vortexed. Samples were gently rocked for 20 minutes and centrifuged at 14000 × *g* for 15 minutes. The top aqueous phase was used for further extraction using GeneJet Plant Extraction kit according to manufacturer’s instructions (Thermo Scientific, Waltham, MA). Fecal samples were extracted using a PureLink™ Microbiome DNA Purification Kit (Thermo Scientific) using a small portion of each fecal sample. All samples, water and fecal, were eluted to 100 µl of elution buffer and stored at −20°C until analysis. The V4 region of the bacterial rRNA gene was selected for community analysis (Parada et al., 2016). Extracted samples were sent to Molecular Research LP (Shallowater, TX) for library prep and amplicon sequencing.

### Microbial community analysis

Raw samples were processed using FASTQ Processor (http://www.mrdnalab.com/16freesoftware/fastq-processor.html) to generate forward, reverse and barcode fastq files for QIIME2 analysis. Pair-end reads were demultiplexed using the q2-demux emp-paired method and denoised using q2-dada2’s denoise-paired method (Bolyen et al., 2018). Sequencing of samples occurred in multiple runs requiring QIIME2 analysis occurring in tandem until after DADA2 analysis, which were then combined for further downstream analysis. Taxonomic composition was generated using a pre-trained Naive Bayes classifier trained on the Greengenes 13_8 99% OTUs trimmed to include the V4 (515F/806R primer pairs). Prior to further downstream analysis, nontarget ASVs, i.e. chloroplast and mitochondria, were removed from sample table and representative sequences.

Constrained analysis of principal coordinates (CAP) was carried out using the vegan R package (Oksanen et al., 2019) to identify factors contributing to differences between sites with respect to environmental factors. Collinearity of environmental factors was examined using a Pearson’s correlation with factors considered collinear with an r^2^ of 0.8 and *P* < 0.05. Weighted Unifrac and Bray-Curtis dissimilarity matrices were used for CAP analysis for environmental samples including total N and P and without total N and P, models were selected by pseudo-AIC using a stepwise algorithm. Alpha and beta diversity, including weighted and unweighted Unifrac principal component analysis, analyses of ASVs was carried out using the phyloseq R package (McMurdie and Holmes, 2013). Dissimlarity between groups was tested using non-parametric PERMANOVA tests with weighted and unweighted Unifrac beta diversity distances (Bik et al., 2016). All statistical analyses were evaluated at an α = 0.05.

An ASV table was exported for use with SourceTracker (Knights et al., 2011) using a parallel version SourceTracker (https://github.com/biota/sourcetracker2) to decrease computational time. Default parameters established in the sourcetracker_for_qiime.py pipeline were used with five runs being conducted. For each sample, the mean proportion (%) for each run were averaged for determination of fecal source contribution.

## Results

### Study Site and Field Samples

A total of 2,261 septic systems, with a density of 94.2 systems per square mile, are located within the Love Creek Watershed, with gravity systems being the dominant system type (Table 2; Fig. 2). Previous analysis by Homsey et al. (Homsey et al., 2015) showed a density of active septic permits of 55.5 permits per square mile, with 1,340 septic systems within the watershed, an increase of approximately 70% of septic systems within the watershed.

**Table 2:** Number of septic systems types within the Love Creek watershed.

| Septic System Type | Number within watershed |
| --- | --- |
| Gravity | 1626 |
| Low Pressure Pipe | 195 |
| Other | 166 |
| Elevated Mound | 94 |
| Pressure Dose | 59 |
| Holding Tank | 35 |
| Alternative | 26 |
| Standard Pressure Dose | 18 |
| Peat | 10 |
| Irrigation | 8 |
| Cesspool | 7 |
| Puraflo Biofilter | 6 |
| Peat Biofilter | 5 |
| Alternative Elevated Sand Mound | 3 |
| Alternative Irrigation | 2 |
| Wisconsin At-Grade | 1 |

**Figure 2.**
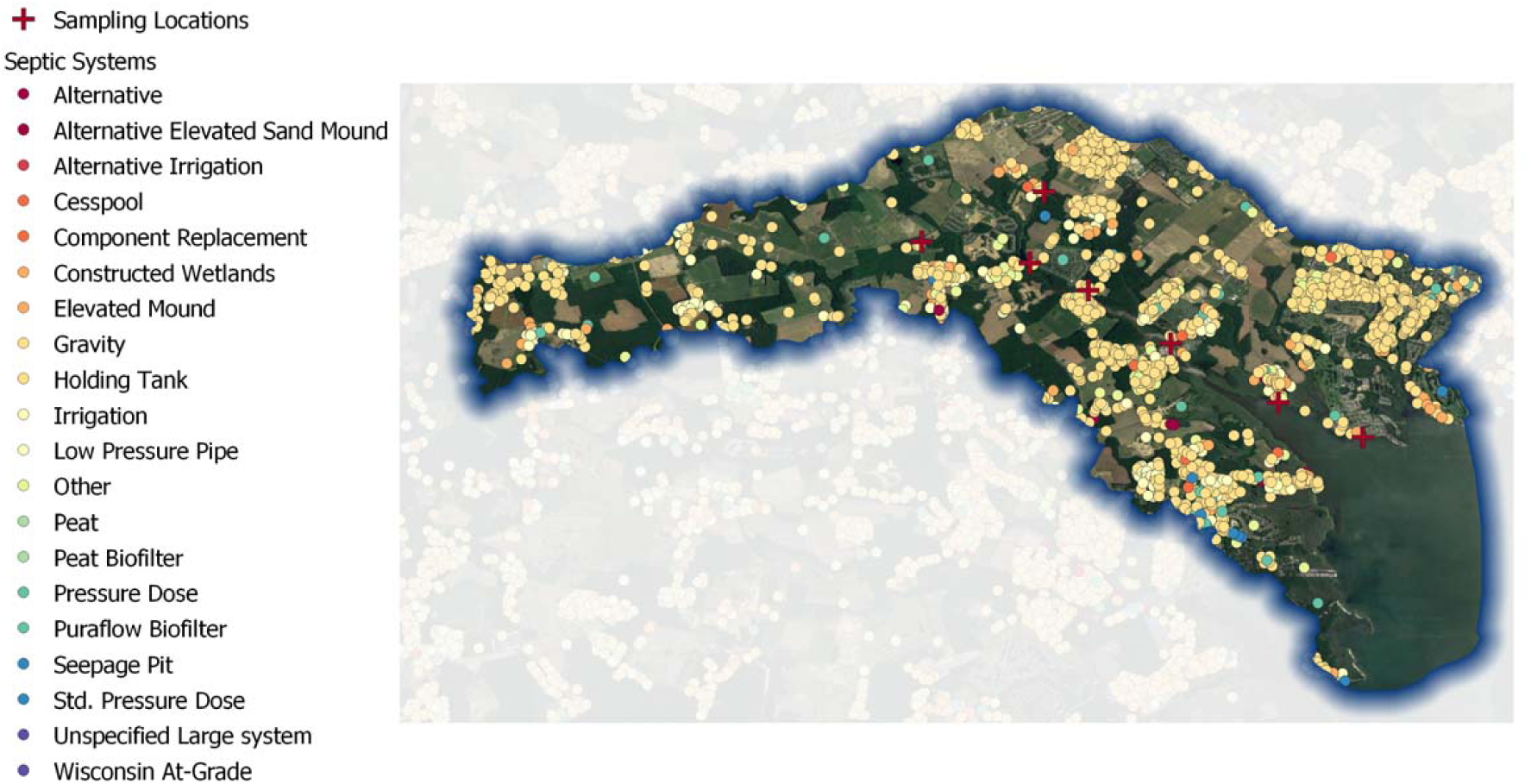
Septic System density in the Love Creek watershed.

Altogether, 74 samples were analyzed from 7 sites within the Love Creek watershed between March and October 2017. Water temperatures during the collection period ranged from 8.3 to 29.7°C, salinity from 0.1 to 41.2, chlorophyll *a* from 0.3 to 382 mg l^−1^, dissolved oxygen from 5.01 to 17.3 mg l^−1^, turbidity from 1 to 44 mg l^−1^, total suspended solids from 1 to 75.9 mg l^−^ 1 and total dissolved solids from 70 to 32,300 mg l^−1^. Organic carbon concentrations ranged from 1.8 to 10.9 mg C l^−1^, dissolved NO_X_ from 0.004 to 7.13 mg N l^−1^, NH_3_ from 0.01 to 0.281 mg N l^−1^, PO_4_ from 0.004 to 0.047 mg P l^−1^, total nitrogen from 0.242 to 7.35 mg N l^−1^, and total phosphorus 0.014 to 0.272 mg P l^−1^. Total *Enterococcus* levels ranged from non-detects to 6,130 mpn 100 ml^−1^ with 27 of 74 samples being above primary contact recreation levels. At Jimtown Road, 5 samples were above PCR, Bundicks Branch 4 samples above PCR, Misty Lane 3 samples above PCR, Route 24 11 of 24 samples above PCR, and West Lane 4 of 9 samples were above PCR. No samples for total *Enterococcus* at Goslee Pond and the mouth of Love Creek were above PCR levels (Fig. 3).

**Figure 3.**
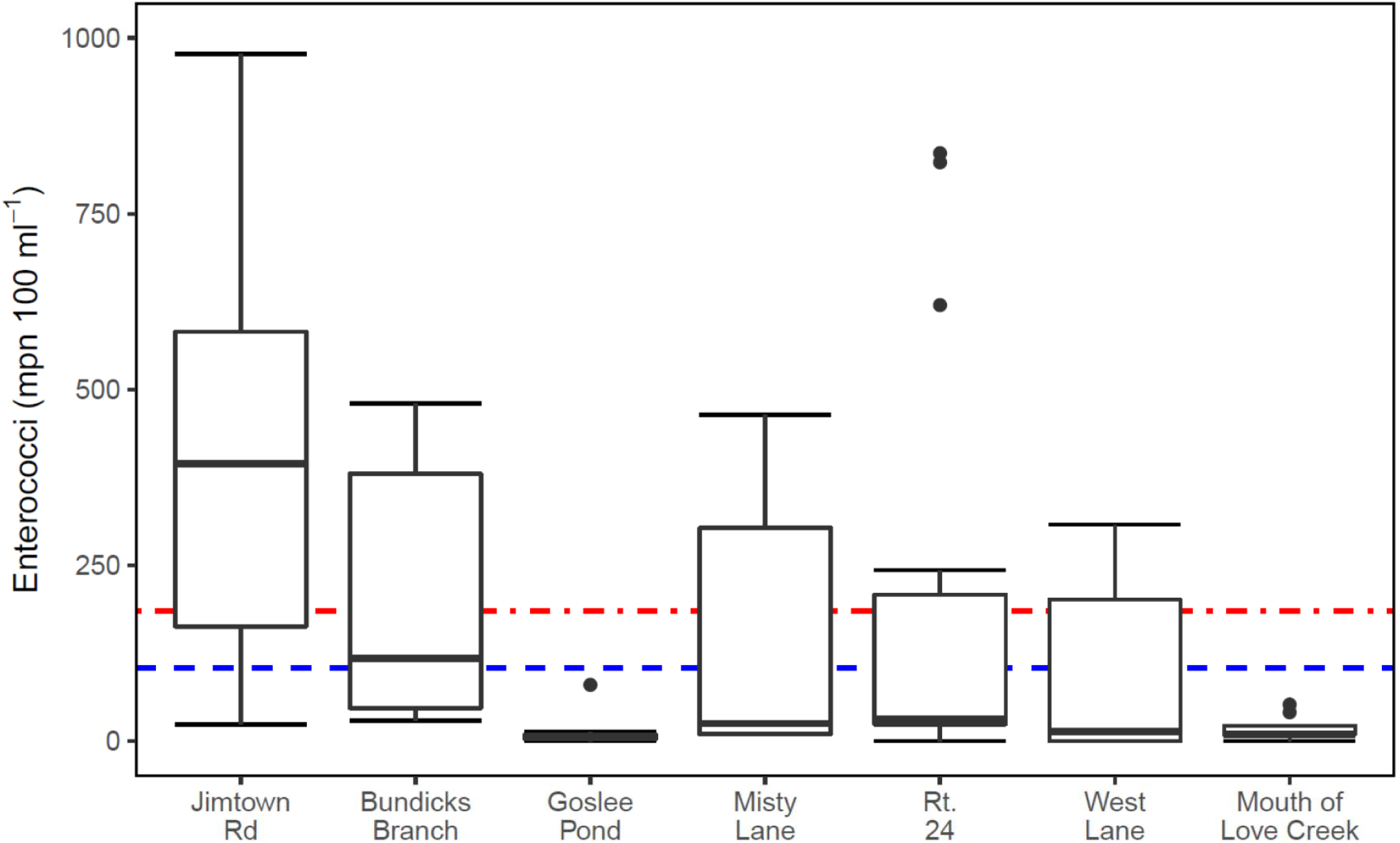
*Enterococcus* levels as determined by Enterolert for each sampling site over the sampling period. PCR for freshwater (blue line) and marine waters (>5 ppt, red line) are also shown.

### Microbial Community Analysis

Taxonomic analysis for 16S data showed that major taxa were largely consistent across sampling sites (Fig. 4). The most abundant phyla across most sample locations were *Proteobacteria, Bacteroidetes*, and *Actinobacteria* (Fig. 4A). *Firmicutes*, one of the most abundant gut microbes (Arumugam et al., 2011), was one of the top phyla for all sampling sites. At Jimtown Rd and Bundicks Branch, *Acidobacteria* and *OP3*, were also in high abundance across all sampling time points. Both the *Acidobacteria* (Kielak et al., 2016) and *OP3* (Arumugam et al., 2011) have been identified from anoxic sediment samples suggesting a potential resuspension of sediments at these sites. At the class level, *Gammaproteobacteria, Alphaproteobacteria, Betaproteobacteria*, and *Flavobacteriia* showed the highest relative abundance across most of the sampling sites (Fig. 4B). For fecal samples at the phylum level, the most abundant phyla were *Firmicutes, Bacteroidetes, Proteobacteria*, and *Euryarchaeota* (Fig. 5A). *Clostridia, Bacilli, Bacteroidia*, and *Methanobacteria* were the dominant classes for fecal samples (Fig. 5B).

**Figure 4.**
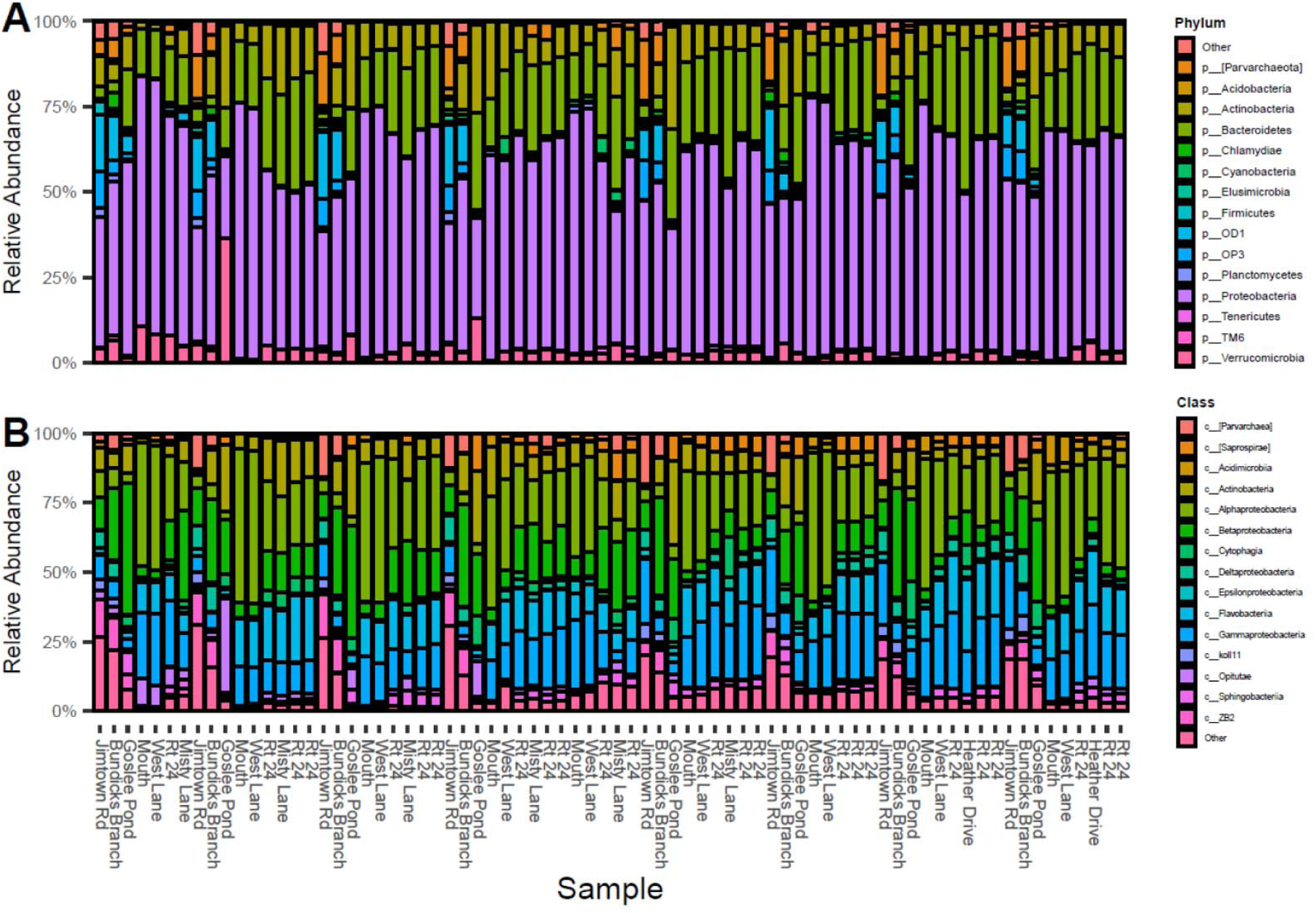
Relative Abundance of bacterial groups at the phylum level (A) and class level (B), showing the top 15 most abundant taxa from monthly samples.

**Figure 5.**
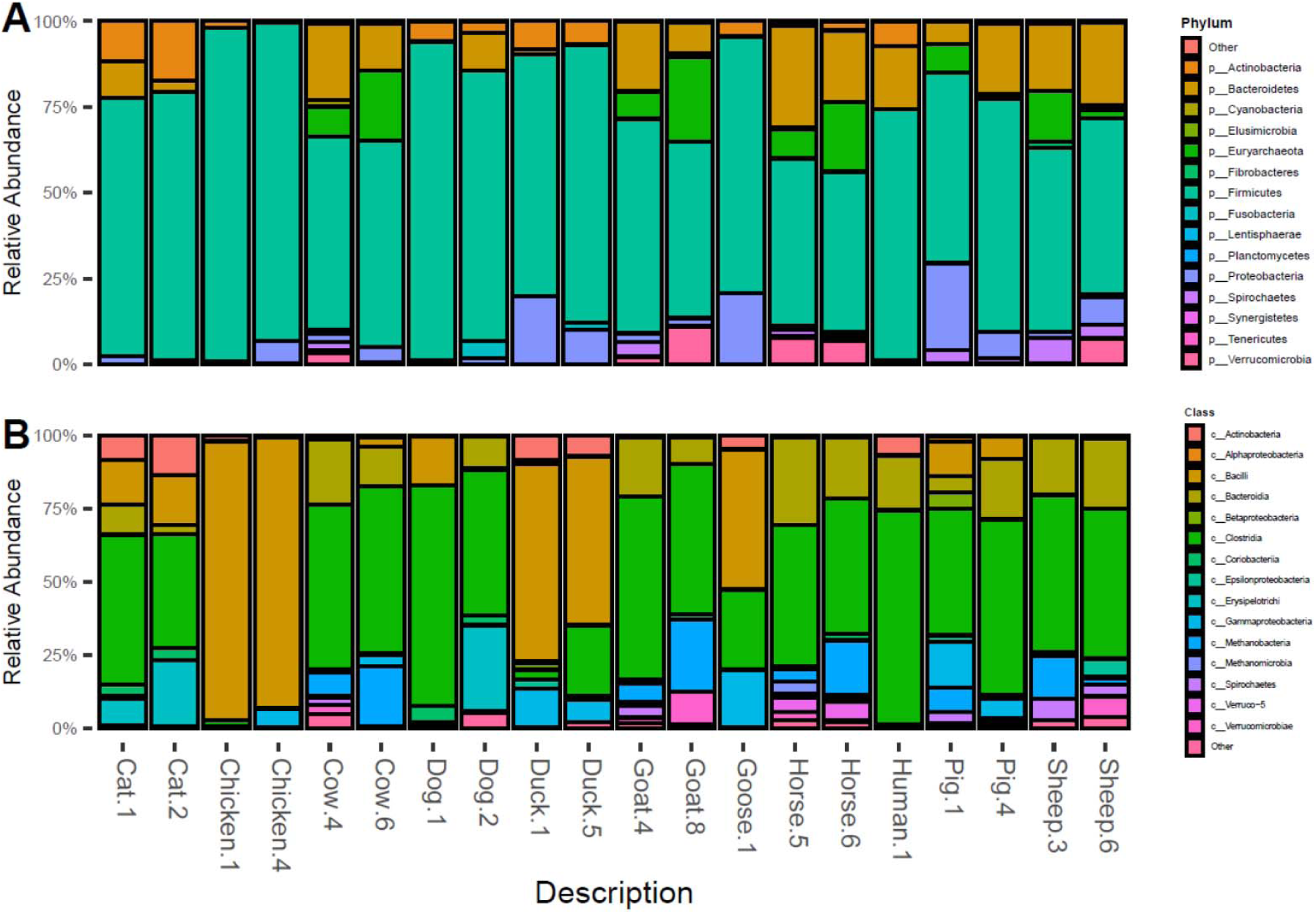
Relative Abundance of bacterial groups at the phylum level (A) and class level (B), showing the top 15 most abundant taxa from fecal samples.

Chlorophyll *a*, dissolved oxygen saturation, NO_X_, specific conductivity, total dissolved solids (TDS) and turbidity were determined to be collinear (Total P, DO, Total N, Salinity, Salinity and total suspended solids, respectively). Previous research has shown a significant correlation to microbial community structure and total dissolved solids (Staley et al., 2014), therefore all CAP evaluations included total dissolved solids. Model selection by stepwise algorithm for weighted Unifrac on all selected factors were TDS, temperature, OP, pH, Salinity, NH_3_, total phosphorus and dissolved oxygen and without total N and P: TDS, temperature, OP, pH, Salinity, NH_3_, chlorophyll *a* and dissolved oxygen (Fig. 6A) with 51.2% of the variation explained on CAP1 and 8.3% explained on CAP2. For Bray-Curtis factors were: salinity, temperature, OP, *Enterococcus*, total N, NH_3_, total organic C, total P, pH, chlorophyll *a*, and dissolved oxygen and without total N and P: salinity, temperature, OP, *Enterococcus*, NO_X_, total organic C, NH_3_, pH, chlorophyll *a*, and dissolved oxygen (Fig. 6B) with 21% of the variation accounted for on CAP1 and 9.7% accounted for on CAP2.

**Figure 6.**
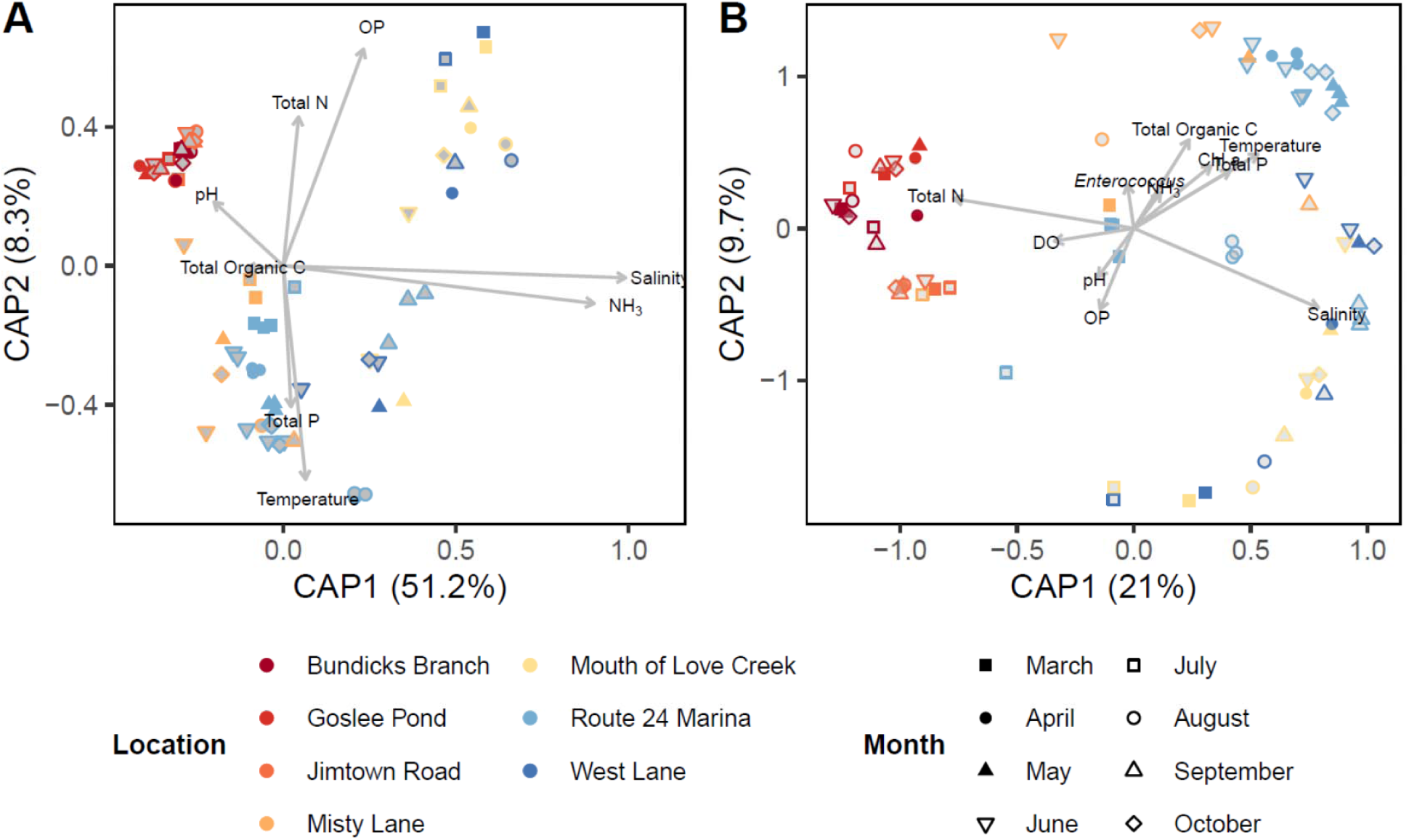
Constrained analysis of principal coordinates showing relationships among species compositions for sites and environmental factors (arrows) for weighted Unifrac (A) and Bray-Curtis (B) for all non-collinearity factors within the watershed (color) and sampling month (shape).

Beta diversity analysis indicated a strong clustering of one distinct group with non-tidal sites (Jimtown Rd, Bundicks Branch and Goslee Pond) separated from tidally driven sites for both weighted Unifrac with 56.9% of the variation accounted for on PCoA1 and 15.7% accounted for on PCoA2 and Bray-Curtis (Fig. 7A and B respectively) with 24.1% of the variation on PCoA1 and 11.6% on PCoA2. Samples collected at the Route 24 marina and Misty Lane clustered together depending on sampling time salinity and temperature (Fig. 6 and 7). PERMANOVA testing showed significant groupings for most environmental factors for both weighted Unifrac and Bray-Curtis (Table 3). Previous research (Bik et al., 2016; Chase et al., 2016) has shown that a putative technical artifact resulting in splitting of 16S sequencing runs. For weighted Unifrac PERMANOVA analysis this artifact was not statistically significant for sample grouping (*P* = 0.0694; Table 3), whereas Bray-Curtis was statistically significant (*P* = 0.0001; Table 3). For fecal samples, the ruminant species clustered together for Bray-Curtis (Fig. 8B) but were more dispersed in weighted Unifrac analysis (Fig. 8A). Additionally, avian samples were clustered in the Bray-Curtis analysis (Fig. 8B) but not weighted Unifrac.

**Table 3:** PERMANOVA test for statistical significance^a^.

| Environmental Factor | Weighted UniFrac *P* Value | Bray-Curtis *P* Value |
| --- | --- | --- |
| Monthly Sampling | **0.0001** | **0.0001** |
| Site Location | **0.0001** | **0.0001** |
| Temperature | **0.0317** | **0.0001** |
| Salinity | **0.0001** | **0.0001** |
| Total dissolved solids | **0.0001** | **0.0001** |
| NH<sub>3</sub> | 0.1908 | **0.0084** |
| Chlorophyll *a* | 0.1305 | **0.0001** |
| Dissolved oxygen | 0.0657 | **0.0057** |
| Total P | 0.1379 | **0.0001** |
| Orthophosphate | **0.0042** | **0.0002** |
| pH | 0.1607 | **0.0159** |
| Total N |  | **0.0001** |
| *Enterococcus* |  | **0.0137** |
| Total organic C |  | **0.0002** |
| NO<sub>x</sub> |  | **0.0001** |
| Sequence Run | 0.0694 | **0.0001** |

<sup>a</sup>Bold values represent $P < 0.05$ .

**Figure 7.**
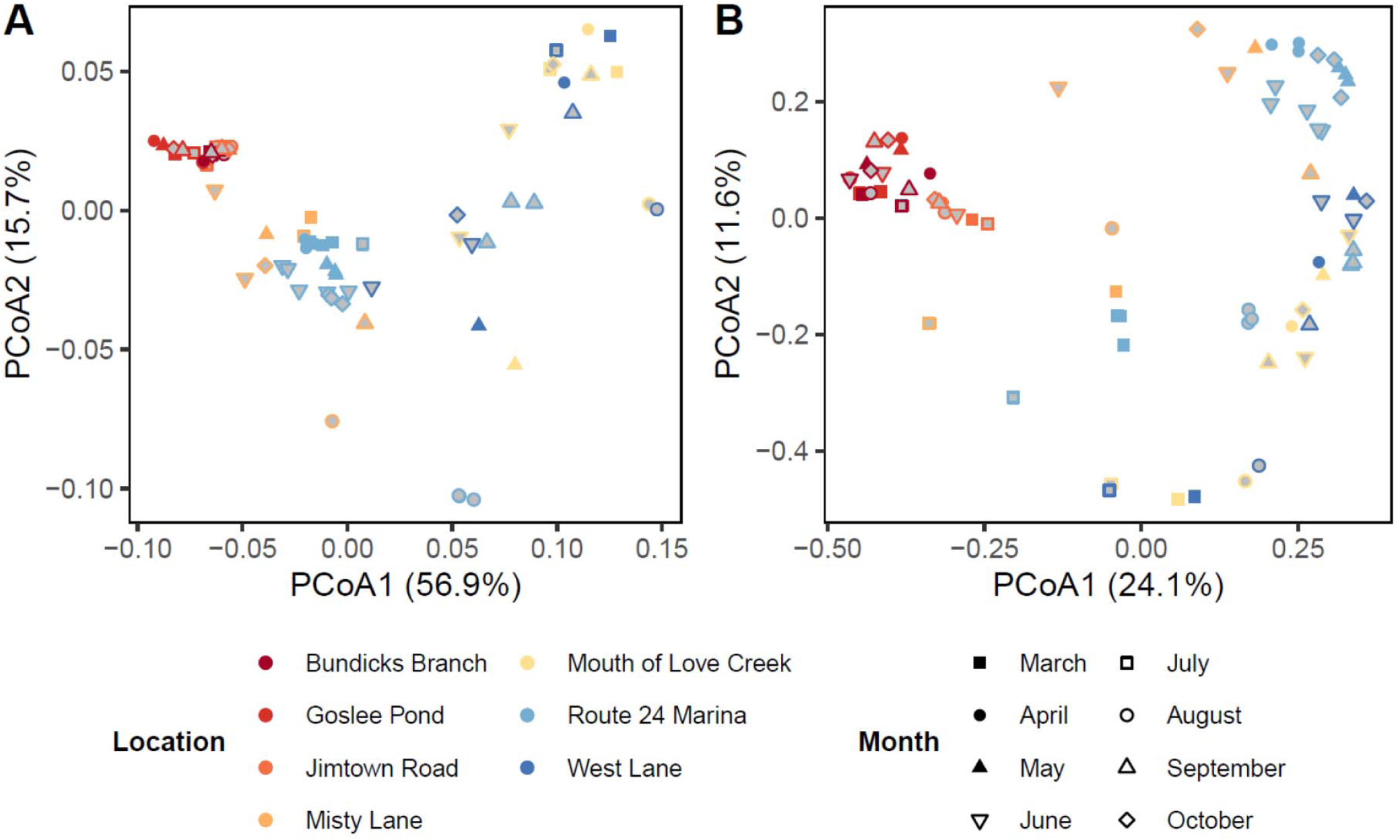
Beta diversity analyses of microbial community for weighted Unifrac PCoA (A) and Bray-Curtis (B) showing sampling location within the watershed (color) and sampling month (shape).

**Figure 8.**
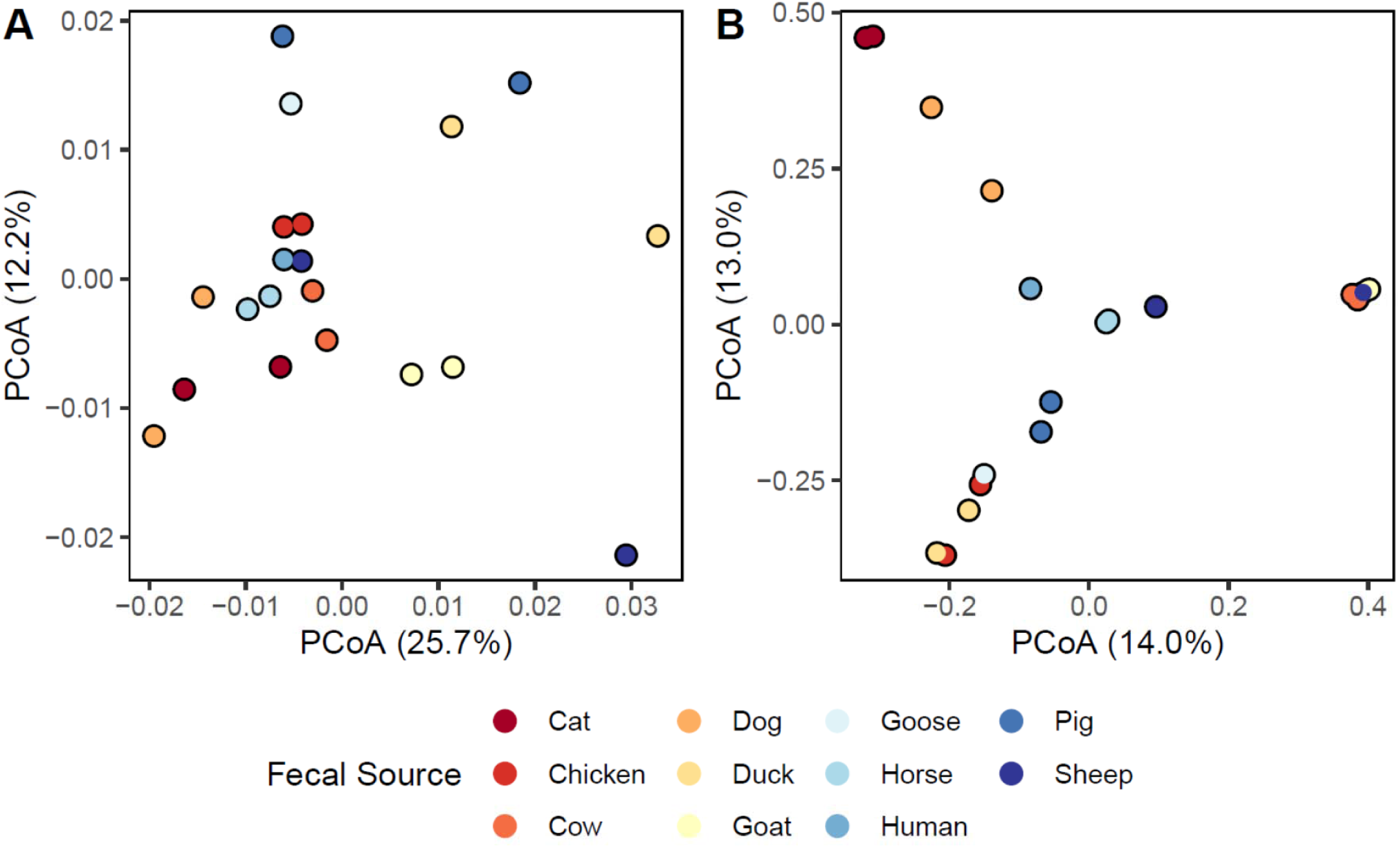
Beta diversity analyses of fecal community for weighted Unifrac PCoA (A) and Bray-Curtis (B) showing fecal samples (color).

### Microbial Sources

SourceTracker analysis on the 16S data indicated that the majority of the bacterial community from all samples were derived from unknown sources (Fig. 9, Fig. 10). For the majority of sites, <40% of the microbial community was assigned to an identified source. At Jimtown Road, the majority of samples were from unknown sources, i.e. wildlife, with samples from April, July, August, September and October above PCR and SCR levels for *Enterococcus* (Fig. 9A). Of those known samples, there was an increase in the amount of cat associated fecal bacteria as the summer progressed. Bundicks Branch had a greater proportion of the microbial community associated with known samples (Fig. 9B) with July through September being above PCR and SCR levels. A decrease of known sources occurred from July to August and from September to October to approximately 5% of the microbial community associated to known sources. Samples collected from Goslee Mill Pond also showed an increase in known sources associated with cat fecal matter (Fig. 9C), however none of the samples were near PCR during the sampling period (Fig. 3).

**Figure 9.**
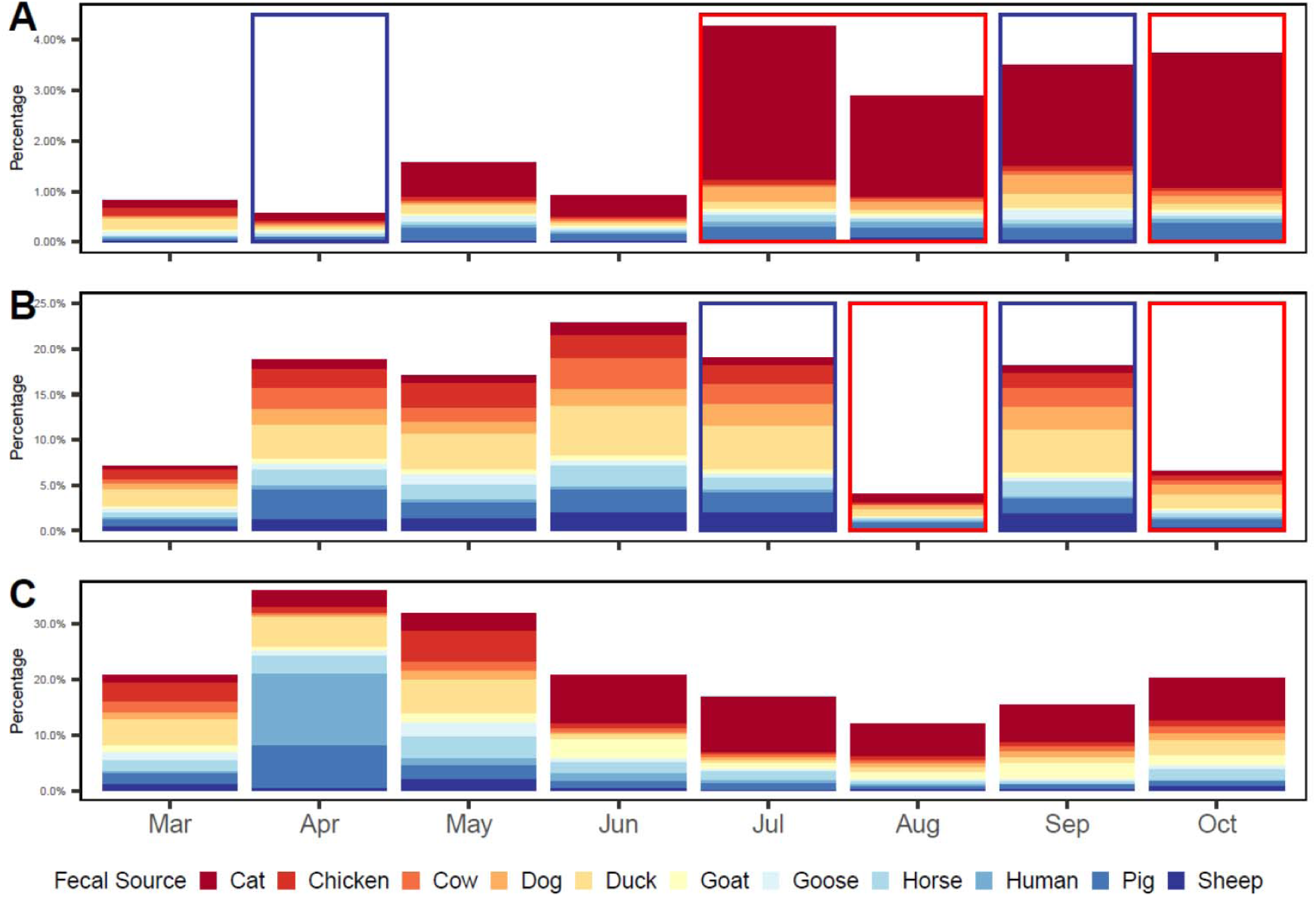
SourceTracker analysis of bacterial assemblages from freshwater sites at Jimtown Road (A), Bundicks Branch (B) and Goslee Mill Pond (C). Sample times that were above PCR are indicated with a blue rectangle and times above both PCR and SCR are indicated with a red rectangle.

**Figure 10.**
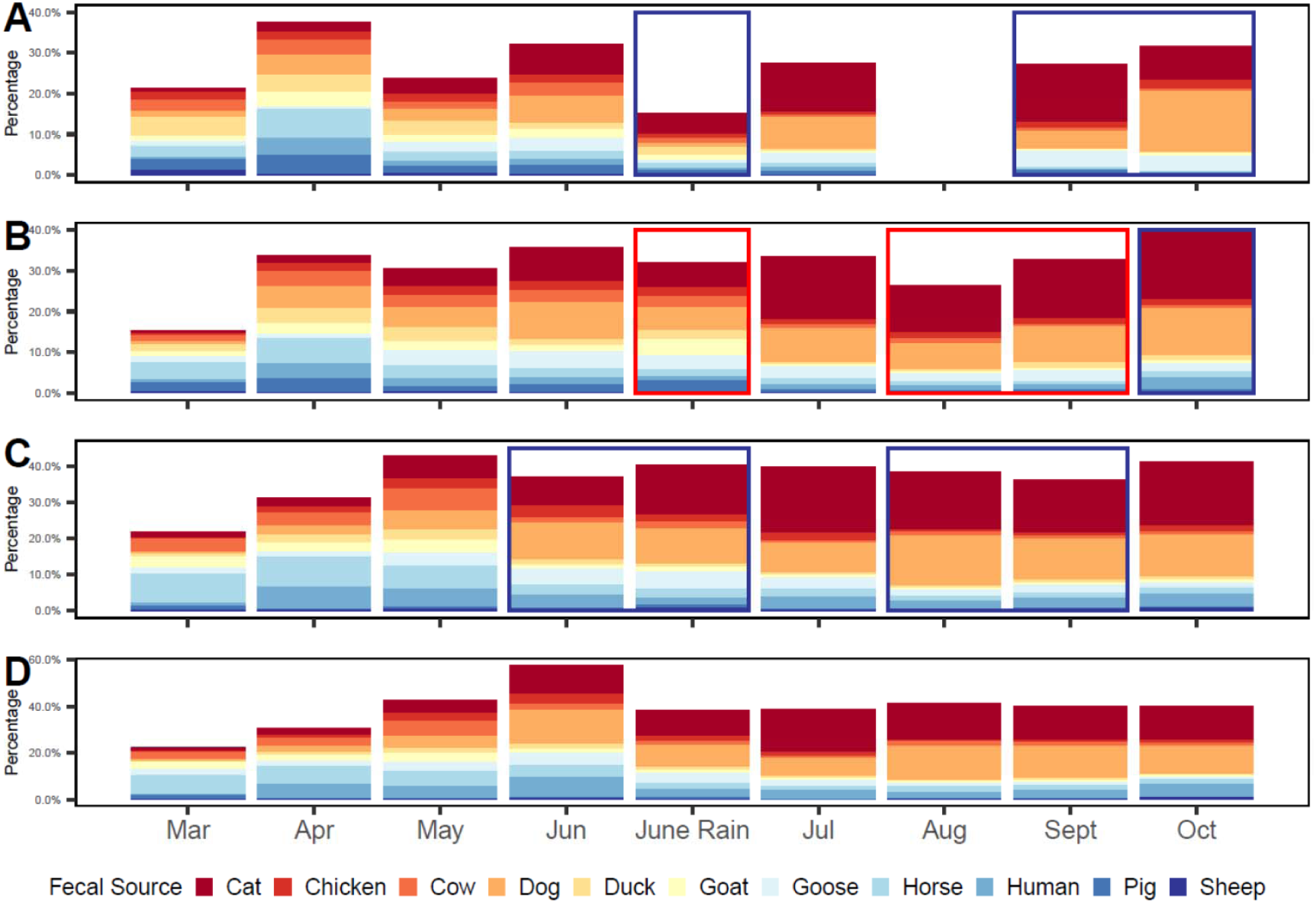
SourceTracker analysis of bacterial assemblages from marine sites at Misty Lane (A), Route 24 marina (B), West Lane (C) and the mouth of Love Creek (D). Sample times that were above PCR are indicated with a blue rectangle and times above both PCR and SCR are indicated with a red rectangle.

Three sampling periods, June rain event, September and October, at Misty Lane were above PCR levels for *Enterococcus* (Fig. 10A). As with Jimtown Road, proportions of the microbial community associated with cat fecal matter increased as the summer progressed. In addition to cat, dog proportions increased over the summer as well, with both of these sources a large portion of the known sources in the microbial community. During the beginning of the sampling period, the housing unit was not occupied, beginning in July onwards the house was occupied with dogs evident on the premises. The Route 24 marina had 4 separate samples above PCR and SCR during the June rain event, and August through October. Both cat and dog sources were a large portion of the known sources at the marina (Fig. 10B). West Lane showed four sampling periods above PCR levels and similar trends in known sources with large proportions of the microbial community associated with dog and cat fecal matter (Fig. 10C). The mouth of Love Creek did not have any samples above contact levels, but did show increases in both cat and dog fecal matter associated microbes as the summer progressed (Fig. 10D).

## Discussion

Current methodologies for determining bacterial contamination in the water bodies of Delaware use testing developed to measure the genus *Enterococcus* via Enterolert©. Primary Contact Recreation (PCR) and Secondary Contact Recreation (SCR) risk-based numeric criteria have been developed by DNREC “to be of non-wildlife origin based on best scientific judgment using available information” (DNREC, 2014a). However, one limitation of the use of *Enterococcus* as a fecal indicator species is the inability to differentiate between fecal sources, i.e. human-related sources and wildlife. Common MST methods use specific bacterial primers from hosts to measure quantities via quantitative real-time PCR (qPCR) (Brown et al., 2017; Harwood et al., 2014). However, the development of primers for fecal sources of interest may be time consuming and previous research has shown potential issues with specificity and sensitivity of qPCR (Brown et al., 2019; Green et al., 2014; Stewart et al., 2013). The usage of NGS methods to define the microbial community of aquatic and fecal samples using amplicon sequence variant (ASV), which can resolve the difference of sequences to a single nucleotide, increases the power of determining potential fecal contamination sources (Callahan et al., 2017; Unno et al., 2018). SourceTracker uses Bayesian models to derive proportions of sources within sink samples, however it has been reported that sources are low concentrations have high variability in estimates (Knights et al., 2011). From a public health standpoint, even at low concentrations of fecal contamination may pose public health risks (Harwood et al., 2014; Henry et al., 2016), thus the high variability in estimates may inhibit detection. Henry et al. (Henry et al., 2016) demonstrated that at low proportions (as low as 0.1%), successful detection by SourceTracker can occur.

Here we evaluated the microbial community and the relationships between environmental and chemical parameters, and the potential usage of next-generation sequencing (NGS) techniques for microbial source tracking (MST) in the Love Creek watershed. The watershed has undergone extensive development over the last several decades resulting in increased the number of potential human-related pollution sources, e.g. septic systems or domestic animals (Fig. 2). In the non-tidal sites of Jimtown Road and Bundicks Branch high bacterial levels occurred during the warmer late summer months (Fig. 9). Both of these sites are highly wooded with low population densities. At Jimtown Road, there is a high density of aging septic systems (Fig. 2), however all samples were below 4% of mapping to known sources suggesting that most of the high bacteria levels are potentially from wildlife sources (Fig. 9A). Two samples at Bundicks Branch were above PCR followed by above SCR the next month. Interestingly, both points above SCR approximately 95% of the community were identified as unknown sources (Fig. 9B). Unlike Jimtown Road, Bundicks Branch the highest proportion comes from duck. In contrast, Goslee Mill pond did not have any high levels of *Enterococcus* during any of the sampling periods suggesting that the pond may act similar to a settling point. Therefore potentially preventing upstream fecal contamination from moving further into the more populated waterways.

Tidally driven sites, with the exception of the mouth of Love Creek, also experienced high *Enterococcus* levels during the warmer months. Land use at Misty Lane is similar to Jimtown Road, with primarily woods with a few smaller vacation homes (Fig. 2). A large proportion of known sources are from both cat and dog for July and September (Fig. 10A). The Route 24 Marina and West Lane demonstrated similar trends for cat and dog and had three periods of *Enterococcus* levels above SCR and one above PCR (Fig. 10B and C respectively). The collection site for Route 24 was directly at a public boat ramp which may increase exposure and health risks for recreational boaters. Additionally, the marina is located within a small development with older septic systems (Fig. 2) which are primarily used during the summer months with few year round residents. West Lane contains more established homes but at a higher density of septic systems (Fig. 2). No high levels of *Enterococcus* were found at the mouth of Love Creek. Sampling occurred at a pier near the edge of the mouth, and may not be indicative of the whole mouth of Love Creek. As with many of the sites within the watershed, cat and dog signals increased during the warmer months, with cat being the largest proportion for both Route 24 and West Lane. The state of Delaware has seen the population of outdoor cats increase over the last several years. As evident by large portions of the known sources from cat signatures (Fig. 9 and 10). Other sources, e.g. goat and horse, are also in high proportions in many of the sites (Fig. 9 and 10). However, observations during collection do not indicate likely locations for these sources, requiring a “boots on the ground” approach for determining likely locations of sources.

Both *Bacteroidetes* and *Firmicutes*, two of the major phyla within human fecal matter (Arumugam et al., 2011; Eckburg et al., 2005), were present in all our samples but not as prevalent as *Proteobacteria*. Other studies have shown that the prevalence of *Proteobacteria* in samples may be an indicator of run-off or storm water (García-Aljaro et al., 2019; Shanks et al., 2013) or from sewage infrastructure, i.e. a transient population (Shanks et al., 2013). SourceTracker analysis indicated that almost all samples from the Love Creek watershed had human signatures from <0.1% to 13%. However, the human signature represents a single individual and does not constitute a representative library. Presently, it is not known how many samples would constitute a representative library (Ahmed et al., 2015) and large variations may occur among human and other fecal samples (Huse et al., 2012). In contrast, Staley, et al. (Staley et al., 2018) suggests that a minimum of 10 individuals are sufficient for accurate fecal contamination detection. However, when few source are used and/or available, results may be better interpreted using broad category classifications, e.g. livestock rather than specific organism. Additionally, Staley, et al. (Staley et al., 2018) demonstrated to increase accuracy of the SourceTracker analysis, geographically associated fecal samples are required. All samples used in this study were geographically located in Delaware. A large library and rarefaction depth may improve identification of sources by SourceTracker (Ahmed et al., 2015). However, previous research has suggested, from a statistical standpoint, rarefying data was inadmissible (McMurdie and Holmes, 2014), as part of our workflow samples were not rarefied.

In this study, we showed that using NGS methods and bacterial community structure can be combined to identify potential sources of fecal pollution in the Love Creek watershed. The use of SourceTracker and NGS are still part of emerging techniques for the tracking of fecal sources. Increasing the library size of known sources, sequencing depth, and greater diversity of sources will require further analysis (Ahmed et al., 2015). Nonetheless, this study is proof that at the state level these methods are capable of giving the start of source tracking. We recommend using these methods as a starting pointing for ground-truthing, as sources may not be evident. One limitation of these methods are the time to develop a library and the computational requirements for large datasets can be a significant challenge. With open-source tools such as QIIME2, SourceTracker and others, are efficient and simple to use, and with the advent of cloud computing large computational clusters are no longer required.

## References

Agnew, L.J., Lyon, S., Gérard-Marchant, P., Collins, V.B., Lembo, A.J., Steenhuis, T.S., Walter, M.T., 2006. Identifying hydrologically sensitive areas: Bridging the gap between science and application. J. Environ. Manage. 78, 63–76. https://doi.org/10.1016/j.jenvman.2005.04.021

Ahmed, W., Staley, C., Sadowsky, M.J., Gyawali, P., Sidhu, J.P.S., Palmer, A., Beale, D.J., Toze, S., 2015. Toolbox Approaches Using Molecular Markers and 16S rRNA Gene Amplicon Data Sets for Identification of Fecal Pollution in Surface Water. Appl. Environ. Microbiol. 81, 7067–7077. https://doi.org/10.1128/AEM.02032-15

American Public Health Association, 2012. Standard methods for the examination of water and wastewater, 22nd ed. American Public Health Association, Washington, DC.

Andres, A.S., 2004. Ground-Water Recharge Potential Mapping In Kent And Sussex Counties, Delaware.

Arumugam, M., Raes, J., Pelletier, E., Le Paslier, D., Yamada, T., Mende, D.R., Fernandes, G.R., Tap, J., Bruls, T., Batto, J.-M., Bertalan, M., Borruel, N., Casellas, F., Fernandez, L., Gautier, L., Hansen, T., Hattori, M., Hayashi, T., Kleerebezem, M., Kurokawa, K., Leclerc, M., Levenez, F., Manichanh, C., Nielsen, H.B., Nielsen, T., Pons, N., Poulain, J., Qin, J., Sicheritz-Ponten, T., Tims, S., Torrents, D., Ugarte, E., Zoetendal, E.G., Wang, J., Guarner, F., Pedersen, O., de Vos, W.M., Brunak, S., Doré, J., Weissenbach, J., Ehrlich, S.D., Bork, P., 2011. Enterotypes of the human gut microbiome. Nature 473, 174–180. https://doi.org/10.1038/nature09944

Bik, H.M., Maritz, J.M., Luong, A., Shin, H., Dominguez-Bello, M.G., Carlton, J.M., 2016. Microbial Community Patterns Associated with Automated Teller Machine Keypads in New York City. mSphere 1, e00226–16. https://doi.org/10.1128/mSphere.00226-16

Bolyen, E., Rideout, J.R., Dillon, M.R., Bokulich, N.A., Abnet, C., Al-Ghalith, G.A., Alexander, H., Alm, E.J., Arumugam, M., Asnicar, F., Bai, Y., Bisanz, J.E., Bittinger, K., Brejnrod, A., Brislawn, C.J., Brown, C.T., Callahan, B.J., Caraballo-Rodríguez, A.M., Chase, J., Cope, E., Da Silva, R., Dorrestein, P.C., Douglas, G.M., Durall, D.M., Duvallet, C., Edwardson, C.F., Ernst, M., Estaki, M., Fouquier, J., Gauglitz, J.M., Gibson, D.L., Gonzalez, A., Gorlick, K., Guo, J., Hillmann, B., Holmes, S., Holste, H., Huttenhower, C., Huttley, G., Janssen, S., Jarmusch, A.K., Jiang, L., Kaehler, B., Kang, K. Bin, Keefe, C.R., Keim, P., Kelley, S.T., Knights, D., Koester, I., Kosciolek, T., Kreps, J., Langille, M.G., Lee, J., Ley, R., Liu, Y.-X., Loftfield, E., Lozupone, C., Maher, M., Marotz, C., Martin, B.D., McDonald, D., McIver, L.J., Melnik, A. V, Metcalf, J.L., Morgan, S.C., Morton, J., Naimey, A.T., Navas-Molina, J.A., Nothias, L.F., Orchanian, S.B., Pearson, T., Peoples, S.L., Petras, D., Preuss, M.L., Pruesse, E., Rasmussen, L.B., Rivers, A., Robeson, II, M.S., Rosenthal, P., Segata, N., Shaffer, M., Shiffer, A., Sinha, R., Song, S.J., Spear, J.R., Swafford, A.D., Thompson, L.R., Torres, P.J., Trinh, P., Tripathi, A., Turnbaugh, P.J., Ul-Hasan, S., van der Hooft, J.J., Vargas, F., Vázquez-Baeza, Y., Vogtmann, E., von Hippel, M., Walters, W., Wan, Y., Wang, M., Warren, J., Weber, K.C., Williamson, C.H., Willis, A.D., Xu, Z.Z., Zaneveld, J.R., Zhang, Y., Zhu, Q., Knight, R., Caporaso, J.G., 2018. QIIME 2: Reproducible, interactive, scalable, and extensible microbiome data science. PeerJ. https://doi.org/ https://doi.org/10.7287/peerj.preprints.27295v2

Brown, C.M., Mathai, P.P., Loesekann, T., Staley, C., Sadowsky, M.J., 2019. Influence of Library Composition on SourceTracker Predictions for Community-Based Microbial Source Tracking. Environ. Sci. Technol. 53, 60–68. https://doi.org/10.1021/acs.est.8b04707

Brown, C.M., Staley, C., Wang, P., Dalzell, B., Chun, C.L., Sadowsky, M.J., 2017. A High-Throughput DNA-Sequencing Approach for Determining Sources of Fecal Bacteria in a Lake Superior Estuary. Environ. Sci. Technol. 51, 8263–8271. https://doi.org/10.1021/acs.est.7b01353

Callahan, B.J., McMurdie, P.J., Holmes, S.P., 2017. Exact sequence variants should replace operational taxonomic units in marker-gene data analysis. ISME J. 11, 2639–2643. https://doi.org/10.1038/ismej.2017.119

Chase, J., Fouquier, J., Zare, M., Sonderegger, D.L., Knight, R., Kelley, S.T., Siegel, J., Caporaso, J.G., 2016. Geography and Location Are the Primary Drivers of Office Microbiome Composition. mSystems 1, 1–18. https://doi.org/10.1128/mSystems.00022-16

Dempster, E.L., Pryor, K. V, Francis, D., Young, J.E., Rogers, H.J., 1999. Rapid DNA extraction from ferns for PCR-based analyses. Biotechniques 27, 66–8.

DNREC, 2014a. Surface Water Quality Standards.

DNREC, 2014b. On-site wastewater permit database [http://WWW.Document]. URL https://data.delaware.gov/Energy-and-Environment/Permitted-Septic-Systems/mv7j-tx3u

Dubinsky, E.A., Butkus, S.R., Andersen, G.L., 2016. Microbial source tracking in impaired watersheds using PhyloChip and machine-learning classification. Water Res. 105, 56–64. https://doi.org/10.1016/j.watres.2016.08.035

Eckburg, P.B., Bik, E.M., Bernstein, C.N., Purdom, E., Dethlefsen, L., Sargent, M., Gill, S.R., Nelson, K.E., Relman, D.A., 2005. Diversity of the human intestinal microbial flora. Science 308, 1635–8. https://doi.org/10.1126/science.1110591

ESRI, 2017. ArcGIS 10.5.

García-Aljaro, C., Blanch, A.R., Campos, C., Jofre, J., Lucena, F., 2019. Pathogens, faecal indicators and human-specific microbial source-tracking markers in sewage. J. Appl. Microbiol. 126, 701–717. https://doi.org/10.1111/jam.14112

Green, H.C., Haugland, R.A., Varma, M., Millen, H.T., Borchardt, M.A., Field, K.G., Walters, W.A., Knight, R., Sivaganesan, M., Kelty, C.A., Shanks, O.C., 2014. Improved HF183 Quantitative Real-Time PCR Assay for Characterization of Human Fecal Pollution in Ambient Surface Water Samples. Appl. Environ. Microbiol. 80, 3086–3094. https://doi.org/10.1128/AEM.04137-13

Harwood, V.J., Staley, C., Badgley, B.D., Borges, K., Korajkic, A., 2014. Microbial source tracking markers for detection of fecal contamination in environmental waters: Relationships between pathogens and human health outcomes. FEMS Microbiol. Rev. 38, 1–40. https://doi.org/10.1111/1574-6976.12031

Henry, R., Schang, C., Coutts, S., Kolotelo, P., Prosser, T., Crosbie, N., Grant, T., Cottam, D., O’Brien, P., Deletic, A., McCarthy, D., 2016. Into the deep: Evaluation of SourceTracker for assessment of faecal contamination of coastal waters. Water Res. 93, 242–253. https://doi.org/10.1016/j.watres.2016.02.029

Homsey, A., Walch, M., Boswell, S., Bason, C., 2015. The State of Your Creek: Love Creek on Rehoboth Bay.

Huse, S.M., Ye, Y., Zhou, Y., Fodor, A.A., 2012. A Core Human Microbiome as Viewed through 16S rRNA Sequence Clusters. PLoS One 7, e34242. https://doi.org/10.1371/journal.pone.0034242

Kielak, A.M., Barreto, C.C., Kowalchuk, G.A., van Veen, J.A., Kuramae, E.E., 2016. The Ecology of Acidobacteria: Moving beyond Genes and Genomes. Front. Microbiol. 7, 1–16. https://doi.org/10.3389/fmicb.2016.00744

Knights, D., Kuczynski, J., Charlson, E.S., Zaneveld, J., Mozer, M.C., Collman, R.G., Bushman, F.D., Knight, R., Kelley, S.T., 2011. Bayesian community-wide culture-independent microbial source tracking. Nat. Methods 8, 761–763. https://doi.org/10.1038/nmeth.1650

Martin, M.J., Andres, A.S., 2008. Analysis and Summary of Water-Table Maps, Report of Investigations No. 73.

McCarthy, D.T., Jovanovic, D., Lintern, A., Teakle, I., Barnes, M., Deletic, A., Coleman, R., Rooney, G., Prosser, T., Coutts, S., Hipsey, M.R., Bruce, L.C., Henry, R., 2017. Source tracking using microbial community fingerprints: Method comparison with hydrodynamic modelling. Water Res. 109, 253–265. https://doi.org/10.1016/j.watres.2016.11.043

McMurdie, P.J., Holmes, S., 2014. Waste Not, Want Not: Why Rarefying Microbiome Data Is Inadmissible. PLoS Comput. Biol. 10, e1003531. https://doi.org/10.1371/journal.pcbi.1003531

McMurdie, P.J., Holmes, S., 2013. phyloseq: An R Package for Reproducible Interactive Analysis and Graphics of Microbiome Census Data. PLoS One 8, e61217. https://doi.org/10.1371/journal.pone.0061217

NRCS, 2016. gSSURGO gridded soil survey geographic database [WWW Document]. URL https://www.nrcs.usda.gov/wps/portal/nrcs/detail/national/home/?cid=nrcs142p2_053628

Oksanen, J., Blanchet, F.G., Friendly, M., Kindt, R., Legendre, P., McGlinn, D., Minchin, P.R., O’Hara, R.B., Simpson, G.L., Solymos, P., Stevens, M.H.H., Szoecs, E.E., Wagner, H., 2019. vegan: Community Ecology Package.

Parada, A.E., Needham, D.M., Fuhrman, J.A., 2016. Every base matters: Assessing small subunit rRNA primers for marine microbiomes with mock communities, time series and global field samples. Environ. Microbiol. 18, 1403–1414. https://doi.org/10.1111/1462-2920.13023

Price, K., 1998. A framework for a Delaware Inland Bays environmental classification. Environ. Monit. Assess. 51, 285–298.

Sallade, Y.E., Sims, J.T., 1997. Phosphorus transformations in the sediments of Delaware’s agricultural drainageways: II. Effect of reducing conditions on phosphorus release. J. Environ. Qual. 26, 1579. https://doi.org/10.2134/jeq1997.00472425002600060018x

Shanks, O.C., Newton, R.J., Kelty, C.A., Huse, S.M., Sogin, M.L., McLellan, S.L., 2013. Comparison of the Microbial Community Structures of Untreated Wastewaters from Different Geographic Locales. Appl. Environ. Microbiol. 79, 2906–2913. https://doi.org/10.1128/AEM.03448-12

Staley, C., Gould, T.J., Wang, P., Phillips, J., Cotner, J.B., Sadowsky, M.J., 2015. Species sorting and seasonal dynamics primarily shape bacterial communities in the Upper Mississippi River. Sci. Total Environ. 505, 435–445. https://doi.org/10.1016/j.scitotenv.2014.10.012

Staley, C., Gould, T.J., Wang, P., Phillips, J., Cotner, J.B., Sadowsky, M.J., 2014. Bacterial community structure is indicative of chemical inputs in the Upper Mississippi River. Front. Microbiol. 5, 1–13. https://doi.org/10.3389/fmicb.2014.00524

Staley, C., Kaiser, T., Lobos, A., Ahmed, W., Harwood, V.J., Brown, C.M., Sadowsky, M.J., 2018. Application of SourceTracker for Accurate Identification of Fecal Pollution in Recreational Freshwater: A Double-Blinded Study. Environ. Sci. Technol. 52, 4207–4217. https://doi.org/10.1021/acs.est.7b05401

Stewart, J.R., Boehm, A.B., Dubinsky, E.A., Fong, T.T., Goodwin, K.D., Griffith, J.F., Noble, R.T., Shanks, O.C., Vijayavel, K., Weisberg, S.B., 2013. Recommendations following a multi-laboratory comparison of microbial source tracking methods. Water Res. 47, 6829–6838. https://doi.org/10.1016/j.watres.2013.04.063

Tarboton, D.G., 1997. A new method for the determination of flow directions and upslope areas in grid digital elevation models. Water Resour. Res. 33, 309–319. https://doi.org/10.1029/96WR03137

Unno, T., Staley, C., Brown, C.M., Han, D., Sadowsky, M.J., Hur, H.-G., 2018. Fecal pollution: new trends and challenges in microbial source tracking using next-generation sequencing. Environ. Microbiol. 20, 3132–3140. https://doi.org/10.1111/1462-2920.14281

Walter, MT, Walter, MF, Brooks, E.S., Steenhuis, T.S., Boll, J., Weiler, K., 2000. Hydrologically Sensitive Areas: Variable Source Area Hydrology Implications for Water Quality Risk Assessment. J. Soil Water Conserv. 53, 277–284.

